# Diversity of Bathyarchaeia viruses in metagenomes and virus-encoded CRISPR system components

**DOI:** 10.1101/2023.08.24.554615

**Authors:** Changhai Duan, Yang Liu, Ying Liu, Lirui Liu, Mingwei Cai, Rui Zhang, Qinglu Zeng, Eugene V. Koonin, Mart Krupovic, Meng Li

## Abstract

Bathyarchaeia represent a class of archaea common and abundant in sedimentary ecosystems. The virome of Bathyarchaeia so far has not been characterized. Here we report 56 metagenome-assembled genomes of Bathyarchaeia viruses identified in metagenomes from different environments. Gene sharing network and phylogenomic analyses led to the proposal of four virus families, including viruses of the realms *Duplodnaviria* and *Adnaviria*, and archaea-specific spindle-shaped viruses. Genomic analyses uncovered diverse CRISPR elements in these viruses. Viruses of the proposed family ‘*Fuxiviridae*’ harbor an atypical type IV-B CRISPR-Cas system and a Cas4 protein that might interfere with host immunity. Viruses of the family ‘*Chiyouviridae*’ encode a Cas2-like endonuclease and two mini-CRISPR arrays, one with a repeat identical to that in the host CRISPR array, potentially allowing the virus to recruit the host CRISPR adaptation machinery to acquire spacers that could contribute to competition with other mobile genetic elements or to inhibition of host defenses. These findings present an outline of the Bathyarchaeia virome and offer a glimpse into their counter-defense mechanisms.

## Main

Bathyarchaeia, formerly the Miscellaneous Crenarchaeotal Group (MCG), is an archaeal class widespread in marine and freshwater sediments^1, 2, 3, 4, 5^. The estimated global abundance of Bathyarchaeia reaches up to 2.0-3.9 × 10^28^ cells, representing one of the most abundant groups of archaea on Earth^6^. Genomic analyses suggest that Bathyarchaeia lead an acetyl-CoA-centered heterotrophic lifestyle with the potential for acetogenesis^6^, methane metabolism^7^, and sulfur reduction^8^. Bathyarchaeia also encompass a variety of genes encoding carbohydrate-active enzymes^9^ and thus likely can unitize various carbohydrates and lignin^10^. The diverse metabolic potential of Bathyarchaeia contributes to their predominance in sedimentary environments, rendering them essential players in the global carbon cycle^5, 8, 11^.

Viruses, as the most abundant biological agents on the planet^12, 13^, reshape the microbial communities through cell lysis^14, 15, 16, 17^ and alter host metabolism via virus-encoded auxiliary metabolic genes (AMGs) ^18, 19, 20^. Bacteria and archaea evolved enormously versatile repertoires of antivirus immune systems. In particular, nearly all archaea and many bacteria encode CRISPR-Cas, the prokaryotic adaptive immunity systems^21^. The CRISPR-Cas system selectively acquires foreign DNA fragments (protospacers) and stores them as spacers in the CRISPR array which is expressed to produce CRISPR (cr) RNAs that serve as guides recognizing the DNA or RNA target and recruiting CRISPR effector nucleases^22^. The CRISPR-Cas systems provide the most reliable basis for viral host prediction by linking host spacers to the cognate virus protospacers^23^.

In response to the host defenses, viruses infecting bacteria and archaea evolved a broad repertoire of counter-defense mechanisms to evade immunity^24, 25, 26^, engaging in the perennial arms race. In particular, many viruses encode diverse anti-CRISPR (Acr) proteins that specifically target different CRISPR-Cas subtypes^27, 28^.

To date, only a limited number of archaeal viruses have been isolated by traditional cultivation-dependent methods. In recent years, an increasing diversity of archaeal viruses has been uncovered through metagenomic data mining^29^, including viruses associated with Asgard archaea^30, 31, 32^, methanotrophic ANME-1 archaea^33^, ammonia-oxidizing archaea^34, 35, 36, 37^ and marine group II Euryarchaeota (Poseidoniales)^38^. However, no viruses linked to Bathyarchaeia, the most prominent group of archaea in the sedimentary ecosystems, have been reported. Here we describe the results of metagenomic analysis revealing a substantial diversity of viruses associated with Bathyarchaeia, some of which encode CRISPR-Cas system components and predicted Acr.

### Discovery of viruses associated with Bathyarchaeia

In total, 367 metagenome-assembled genomes (MAGs) of Bathyarchaeia, including 68 high-quality MAGs (Extended Data Table S1), were obtained from our previous results^8, 39^ and public databases. Phylogenetic analysis based on a concatenated alignment of an optimized set of 51 marker proteins (see Methods) yielded 8 major clades of Bathyarchaeia that correspond to the orders in the latest taxonomy^40^ (Fig. 1a and Extended Data Fig. S1).

**Fig. 1.**
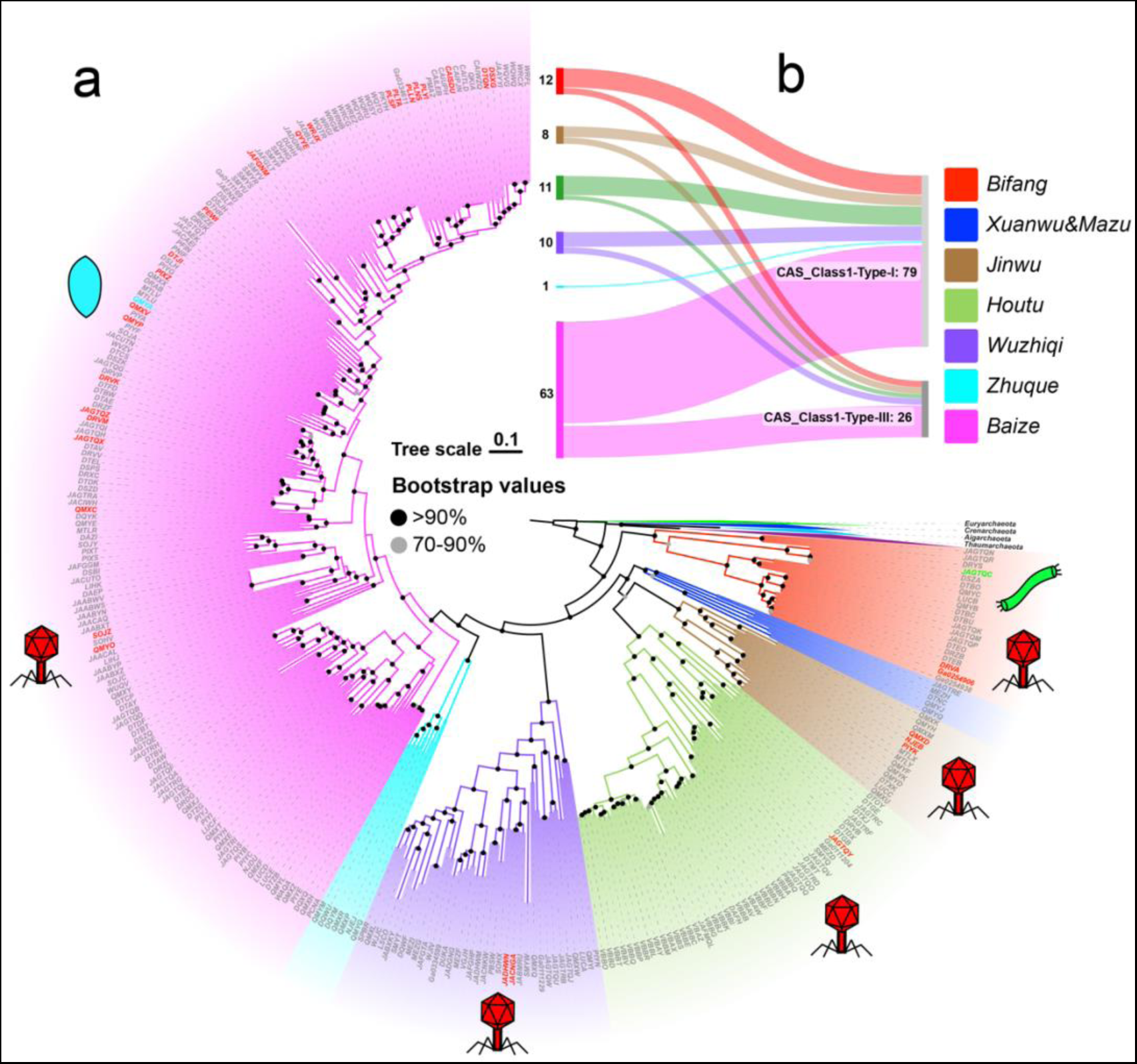
Distribution of identified viruses across the evolutionary tree of the Bathyarchaeia. **a**) Maximum likelihood tree of Bathyarchaeia was reconstructed based on the modified set of 51 marker genes. The outgroup was taken from reference^102^ and includes representatives of the phyla Euryarchaeota, Crenarchaeota, Aigarchaeota and Thaumarchaeota.. Bathyarchaeia orders are highlighted in different background colors. Viruses identified in this study are indicated with colored symbols denoting the respective virion architectures, and their corresponding hosts are highlighted with the same color. Red icosahedron represents viruses in the realm *Duplodnaviria*, green rod represents viruses in the realm *Adnaviria*, and blue spindle represents the spindle-shaped virus. **b**) Subtypes of CRISPR-Cas systems distributed in different Bathyarchaeia orders. Bifang, Bifangarchaeales; Xuanwu, Xuanwuarculales; Jinwu, Jinwuousiales; Mazu, Mazuousiales; Houtu, Houtuarculales; Wuzhiqi, Wuzhiqiibiales; Zhuque, Zhuquarculales; Baize, Baizomonadales.

We then searched for CRISPR-Cas systems in Bathyarchaeia MAGs and identified type I and type III CRISPR-*cas* loci in over one-fifth of the MAGs. Typically, a bathyarchaeal genome carries a single CRISPR-Cas system of, either type I (53 high quality MAGs) or type III (15 high quality MAGs). Notably, however, 11 MAGs were found to encompass both types of CRISPR-Cas systems (Fig. 1b). To identify putative viruses of the Bathyarchaeia, a dataset of CRISPR spacers was compiled from all collected MAGs. After automatic screening followed by manual inspection (see Methods), 49 high-confidence CRISPR array from different Bathyarchaeia orders were identified, containing 1,602 spacers (Extended Data Fig. S2). The spacers were used to search for protospacers in the viral sequences of IMG/VR v3 database^41^. Additionally, a search for potential proviruses in the host genomes was conducted (see Methods). In total, 56 contigs were assigned to Bathyarchaeia viruses based on CRISPR spacer matches or unequivocal integration into the host genome. Among these, 54 contigs were found to encode major capsid proteins (MCPs), attesting to their viral nature^42^. Six viral contigs corresponded to complete genomes, as indicated by the presence of terminal repeats (Extended Data Table S3).

### Four putative families of Bathyarchaeia viruses

Protein sharing network analysis identified four distinct groups of Bathyarchaeia viruses, including a large group that consisted of 6 viral clusters (VCs) (Fig. 2). Additionally, we carried out a genome-wide sequence similarity comparison and examined the phylogenies of hallmark genes in Bathyarchaeia viruses, comparing them with known archaeal viruses (Extended Data Fig. S3). Based on the results of these analyses, we identified four putative family-level groups of Bathyarchaeia viruses. The families ‘*Fuxiviridae*’ and ‘*Kunpengviridae*’ include head and tail viruses of the class *Caudoviricetes* in the realm *Duplodnaviria*. The family ‘*Chiyouviridae*’ consists of filamentous viruses of the archaea-specific realm *Adnaviria*^43^. The fourth putative family, ‘*Huangdiviridae*’, with only one representative genome, includes an archaea-specific spindle-shaped virus; the spindle-shaped viruses have not yet been classified at higher taxonomy ranks (Fig. 2).

**Fig. 2.**
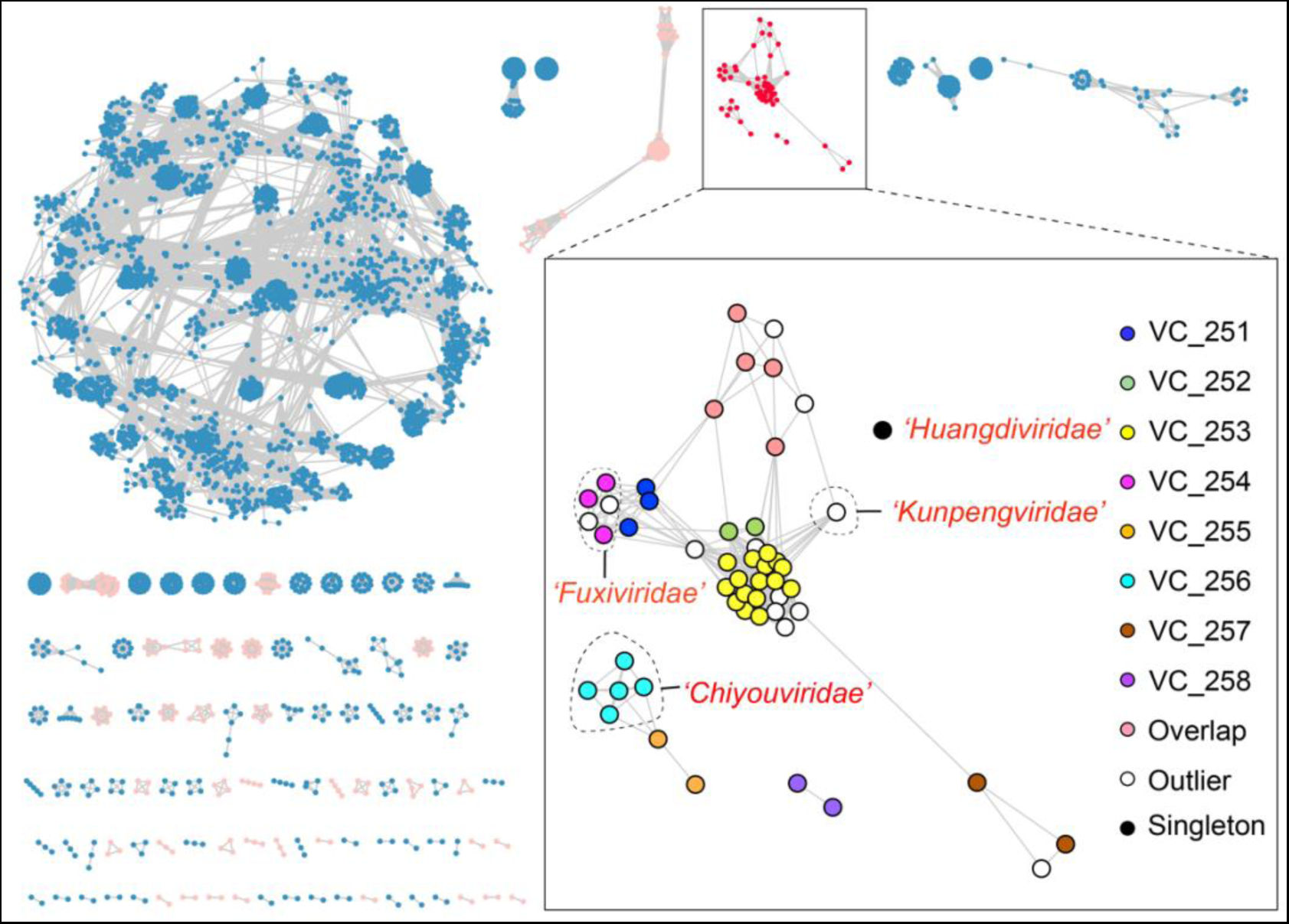
Classification of Bathyarchaeia viruses based on the whole-genome protein-sharing network with other prokaryotic viruses. The whole-genome protein-sharing network analysis was constructed using vConTACT2 v.0.11.3 for the taxonomic assignment of 56 Bathyarchaeia viral genomes. Bathyarchaeia viruses clusters are highlighted in red in the complete network. Bathyarchaeia viruses are assigned to 4 distinct groups, including one large cluster. Viral clusters (VCs) are indicated with differently color circles. The proposed virus families including complete genomes are separated by dashed lines and appended with the corresponding names in red. Archaeal viruses are in pink, and bacterial viruses are in blue. The networks were visualized with Cytoscape v.3.9.1.

The proposed family ‘*Fuxiviridae*’ is represented by three nearly identical complete genomes (Fuxivirus) that encompass protospacers targeted by type I-A CRISPR spacers from the Bathyarchaeia order Bifangarchaeales (Fig. 1 and Fig. 3). Fuxivirus has a smaller genome compared to the typical size of archaeal viruses of the class *Caudoviricetes* (median size of 54.3 kb, n=44), with a length of 31,982 bp (Fig. 3a). Fuxivirus encodes all the hallmark proteins of *Caudoviricetes*, namely, a HK97-like MCP (Extended Data Fig. S3b and Fuxivirus_34 in Extended Data Table S4), a portal protein (Fuxivirus_28), a terminase large subunit (Fuxivirus_27), a tail tube protein (Fuxivirus_38) and several other tail components (Fig. 3a and Extended Data Table S4) which are similar to previously characterized archaeal tailed viruses^44^. In addition to the viral hallmark genes, Fuxivirus encodes several putative DNA-binding proteins, such as predicted transcription factors containing Zn-finger and winged-helix-like domains (Fig. 3a and Extended Data Table S4) that likely regulate viral gene expression^45^. Notably, Fuxivirus also encodes CRISPR-Cas system components, namely, a type IV *cas* gene cluster (Fuxivirus_1 to Fuxivirus_5 in Extended Data Table S4) with a mini CRISPR array, and a Cas4 protein (Fuxivirus_12) (Fig. 3a and see below).

**Fig. 3.**
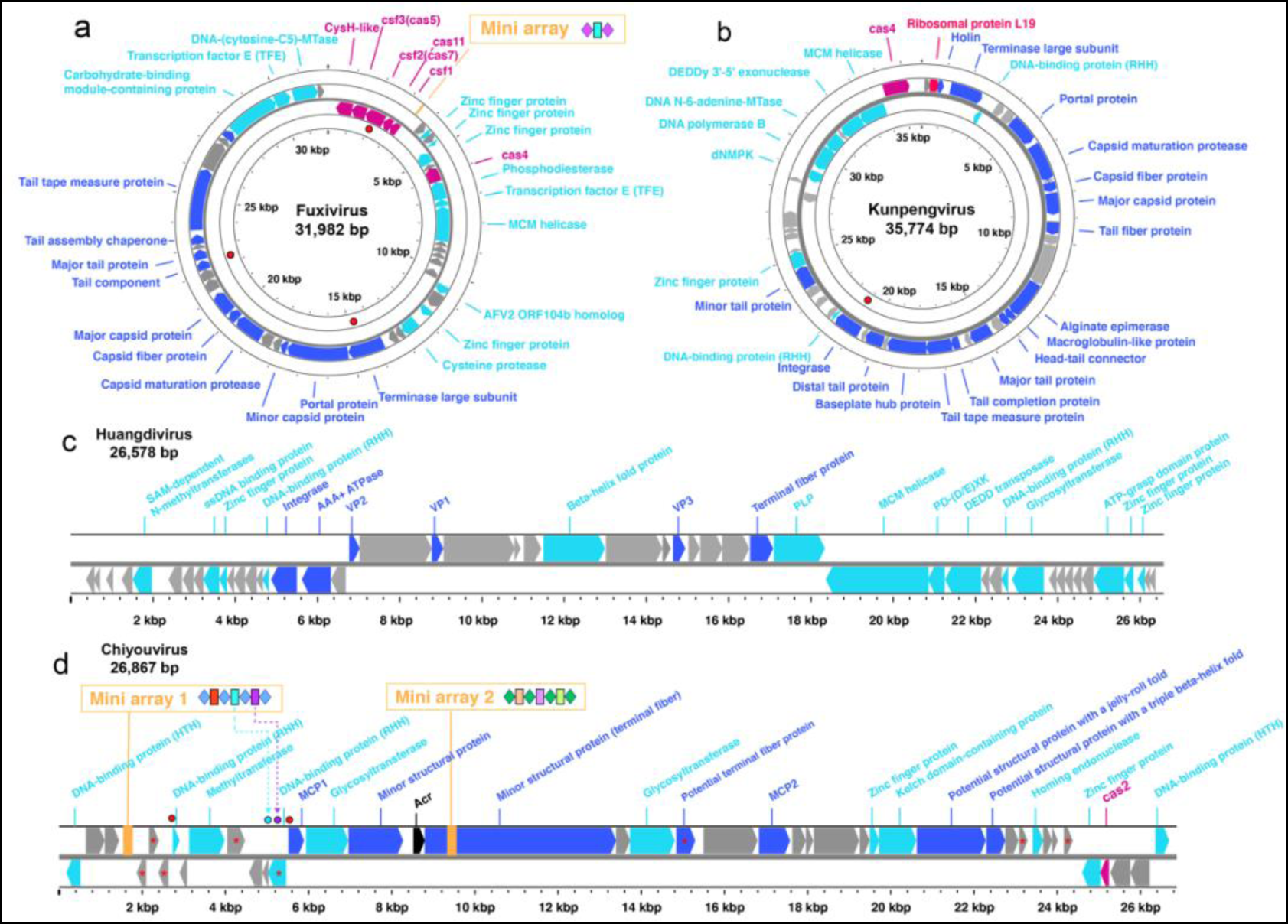
Maps of complete genomes of Bathyarchaeia viruses. **a**) Genome map of Fuxivirus. **b**) Genome map of Kunpengvirus. **c**) Genome map of Huangdivirus. **d**) Genome map of Chiyouvirus. Genes annotated by HHblits with a probability greater than 95% are shown as different colors. Genes related to type IV B CRISPR-Cas system, Cas4 and Cas2 are indicated in rose pink, mini-CRISPR array in vibrant orange, genes specific to viruses in dark blue, predicted Acr in black, other annotated genes in light blue. The positions of targeted protospacers are indicated with red circle. The organization of CRISPR mini-arrays is shown above the genome maps. In Chiyouvirus CRISPR array 1, the self-targeting spacers are highlighted in light blue and purple. Their corresponding target sites on the genome are marked with circles in the same color. Transmembrane proteins of Chiyouvirus predicted using CCTOP^103^ server are indicated with red asterisks. Detailed gene annotations are in Supplementary table S4.

The proposed family ‘*Kunpengviridae*’ includes one complete viral genome (Kunpengvirus), that is targeted by a single spacer (100% match) from *Bathyarchaeia* sp. QMXD of the order Jinwuousiales (Fig. 1). (Fig. 3b). Kunpengvirus encodes the hallmark proteins of *Caudoviricetes*, as well as a suite of tail proteins including the baseplate protein (Fig. 3b and Extended Data Table S4). Kunpengvirus also encodes an integrase (Kenpengvirus_28), suggesting that it can integrate into the host genome as a provirus. In addition, this virus encompasses genes for a deoxynucleoside monophosphate kinase (dNMPK) (Kenpengvirus_41), an MCM-like helicase (Kenpengvirus_46) and a family B DNA polymerase (Kenpengvirus_42) (Fig. 3b and Extended Data Table S4) which are indicative of (at least, partially) autonomous genome replication^46^. Notably, unlike in other tailed viruses, the proofreading DEDDy 3’-5’ exonuclease (Kenpengvirus_45) and the family B DNA polymerase are encoded by two distinct genes. Additionally, Kunpengvirus encodes a homolog of ribosomal protein bL19 (Kunpengvirus_2 in Extended Data Table S4), which is typically present in bacteria and eukaryotes (chloroplasts and mitochondria) but not in archaea, except for *Candidatus* Aenigmarchaeota. The bL19 is located at the 30S-50S ribosomal subunit interface and is thought to contribute to the structure and function of the aminoacyl-tRNA binding site^47^. A number of ribosomal proteins, including bL19, were previously identified in bacterial viruses^48^, but the only ribosomal protein so far detected in archaeal viruses had been L21e^44^. The presence of bL19 in Kunpengvirus suggests that modification of the host translation apparatus by archaeal viruses is more common than currently recognized. Similar to the Fuxivirus, Kunpengvirus also encodes a Cas4 endonuclease (Fig. 3b and Kenpengvirus_47 in Extended Data Table S4).

A unique viral genome (Huangdivirus) representing the proposed family ‘*Huangdiviridae*’ was identified as an apparent provirus in *Bathyarchaeia* sp. QMYA of the order Baizomonadales from a deep-sea hydrothermal vent (Fig. 1). Sensitive sequence comparison using HHsearch identified three virus-encoded structural proteins (VPs) VP1-3 (Fig. 3c and Extended Data Table S4), homologous to the structural proteins of archaeal spindle-shaped viruses of the family *Fuselloviridae*, as well as an AAA+ ATPase (Huangdivirus_17, HHblits best hit to ATV ATPase, with 99.8% probability). Structural predictions for VP1 (Huangdivirus_21) and VP3 (Huangdivirus_28) indicated that both proteins contain two hydrophobic α-helices connected by a short turn (Extended Data Fig. S5a) resembling the typical structure of the MCPs of spindle-shaped viruses^49^. Huangdivirus VP2 (Huangdivirus_19) is most closely related to the viral DNA-binding protein VP2 of Sulfolobus spindle-shaped virus (SSV1)^50^ (HHblits probability of 99.49%). As in the case of SSV1^50^, we detected the consensus glycosylation motifs (N-X-S/T) in Huangdivirus VP1 and VP3 which may be glycosylated by the virus-encoded glycosyltransferase (Huangdivirus_40 and Extended Data Fig. S4b). Capsid protein glycosylation is thought to contribute to the virion stability, particularly for viruses infecting hyperthermophilic hosts^50^. Similar to fuselloviruses infecting hyperthermophilic archaea of the order Sulfolobales^51^, Huangdivirus encodes a putative integrase of the tyrosine recombinase superfamily (Fig. 3c and Huangdivirus_16 in Extended Data Table S4) which is likely to be responsible for viral DNA integration into the host chromosome. Notably, Huangdivirus encodes a predicted polysaccharide lyase (Fig.3c and Huangdivirus_33 in Extended Data Table S4) that might be involved in the break-down of the S-layer polysaccharides^52, 53^ of host cell wall during infection.

‘*Chiyouviridae*’ is a potential new family of filamentous viruses in the order *Ligamenvirale* (class *Tokiviricetes*, realm *Adnaviria*). ‘*Chiyouviridae*’ is represented by one complete viral genome (Chiyouvirus) that is targeted by two spacers of *Bathyarchaeia* sp. JAGTQC in the order Bifangarchaeales (Fig. 1 and Fig. 3d). Phylogenomic analysis of all available *Tokiviricetes*^54^ genomes recapitulated the previously established relationships and showed that Chiyouvirus forms a separate clade within the order *Ligamenvirales*, most closely related to the families *Rudiviridae* and *Ungulaviridae* (Extended Data Fig. S6a). Whole proteome comparison showed less than 50% average amino acid identity (AAI) between protein homologs from Chiyouvirus and members of other viral families, with the highest AAI (46%) with the genus *Icerudivirus* of the family *Rudiviridae* (Extended Data Fig. S6b). Similar to all members of the families *Ungulaviridae* and *Lipothrixviridae* but only some members of the family *Rudiviridae*^54^, Chiyouvirus encodes two MCPs (Chiyouvirus_14 and Chiyouvirus_23), each comprising an alpha-helix bundle (Fig. 3d and Extended Data Fig. S6c). In rudiviruses, unlike other members of the realm *Adnaviria*, the second MCP paralog, when present, is not incorporated into virions^55^. Thus, it remains unclear whether both Chiyouvirus MCPs are involved in virion formation. In addition, Chiyouvirus encodes a large minor structural protein (Fig. 3d and Chiyouvirus_18 in Extended Data Table S4), a homolog of SIRV2 P1070, which is thought to be involved in the formation of the virion terminal filaments that are responsible for host recognition^56, 57^. Predictions for transmembrane proteins indicated that Chiyouvirus encodes 8 potential transmembrane proteins (Fig. 3d) suggesting that it is a membrane-enveloped filamentous virus, similar to the adnaviruses in the families *Lipothrixviridae*, *Ungulaviridae* and *Tristromaviridae*^58^.

### CRISPR-Cas systems and potential counter defense mechanisms in Bathyarchaeia viruses

We explored the type IV CRISPR-Cas system encoded by ‘*Fuxiviridae*’ in greater detail, including phylogenetic analysis and gene locus comparison. The viral CRISPR-*cas* locus encodes all typical components of the type IV CRISPR-Cas effector module (Fig. 3a and Extended Data Table S4), in particular, the signature protein Csf1 (Fuxivirus_5 in Extended Data Table S4), the apparent large subunit of the effector complex, but no adaptation module. Phylogenetic analysis of Cas proteins, including Csf2 (Cas7) (Fuxivirus_3 in Extended Data Table S4), the most conserved protein in the type IV systems^59^, showed that CRISPR-Cas system of ‘*Fuxiviridae*’ belongs to the IV-B subtype (Extended Data Fig. S7). Additionally, a CysH-like protein (Fuxivirus_1 in Extended Data Table S4), which is tightly associated with the type IV-B systems, is encoded in the virus genome adjacent to Csf3 (Cas5) (Fuxivirus_2; Fig. 3a). Phylogenetic analysis showed that Fuxivirus CysH-like protein does not belong to the viral CysH branch but is rather associated with CysH-like proteins from other type IV-B systems (Extended Data Fig. S7c). Previously, type IV CRISPR-Cas systems have been observed in plasmids and prophages^60, 61^ as well as several lytic phages^62^, but not in archaeal viruses.

Most type IV-B systems lack both the adaptation module and a CRISPR array, but the type IV-B CRISPR-*cas* locus in the Fuxivirus genomes contains a CRISPR mini-array that consists of two repeats and a spacer (Fig. 3a). A type IV-B system has been reported to form a filamentous RNP complex which predominantly assembles on non-CRISPR RNAs, without apparent sequence specificity, suggesting a function distinct from adaptive immunity^63^. However, given the presence of the mini-array, we hypothesize the Fuxivirus type IV-B complex could incorporate spacers, conceivably, by recruiting the host adaptation machinery, and utilize a virus-encoded crRNA to target host DNA or other coinfecting mobile genetic elements (MGEs), thus, being potentially involved in evading host immunity and/or inter-MGE conflicts^64^. However, no full-length protospacer matches for the Fuxivirus mini-array spacer sequence were found in either the host or other MGEs, so that further validation of the counter-defense function of the Fuxivirus type IV-B CRISPR-Cas system is needed.

The inferred Fuxivirus host genome, *Bathyarchaeia* sp. DRVA, contains two CRISPR arrays and two *cas3* gene but lacks a complete CRISPR-Cas system. Another genome of *Bathyarchaeia* sp. JAGTQM, from the same order Bifangarchaeales, was found to share identical CRISPR repeat and a closely similar Cas3 protein (84.03% identity, 100% coverage) with *Bathyarchaeia* sp. DRVA. However, *Bathyarchaeia* sp. JAGTQM contains complete type I-A and III-D CRISPR-Cas system, suggesting the presence of a CRISPR-Cas system(s) in the DRVA genome as well. Notably, we found that the *cas7* gene of Fuxivirus was targeted by a host spacer (Fig. 3a), suggesting that the virus-encoded type IV-B system, and with it, possibly, the virus reproduction, can be inhibited by the host CRISPR-Cas systems.

The Cas4 homolog encoded by Fuxivirus could be an additional counter-defense factor (Fig. 3a and Fuxivirus_12 in Extended Data Table S4). Cas4 is a P-D/ExK family nuclease that is a common component of CRISPR-Cas systems that assists Cas1-Cas2 integration complexes in the acquisition of CRISPR spacers^65^, but many Cas4 homologs are encoded outside CRISPR-*cas* loci^66^. The Cas4 homologs are encoded by both ‘*Fuxiviridae*’ (Fuxivirus_12) and ‘*Kunpengviridae*’ (Kenpengvirus_47). Phylogenetic analysis showed that these Bathyarchaeia viral Cas4 proteins are most closely related to Cas4 homologs encoded by Sulfolobus-infecting rudiviruses (Extended Data Fig. S7b), suggesting the possibility of horizontal gene transfer between unrelated archaeal viruses. Multiple sequence alignment confirmed that two Cas4 homologs encoded by ‘*Fuxiviridae*’ and ‘*Kunpengviridae*’ share nearly all conserved amino acids with the rudivirus SIRV2-encoded Cas4 homolog and are thus predicted to be active nucleases (Extended Data Fig. S8). Previous studies have demonstrated that overexpression of the SIRV2-encoded Cas4 in the archaeal host substantially reduced the efficiency of exogenous spacers acquisition by the host CRISPR-Cas system^67^. Thus, the Fuxivirus and Kunpengvirus Cas4 might inhibit spacer acquisition by the CRISPR systems of Bathyarchaeia hosts. However, involvement of this nuclease in the virus genome replication cannot be ruled out either^68^.

We identified two mini-arrays in Chiyouvirus genome, each containing 4 repeats and 3 spacers (Fig. 3d), and the repeats in Chiyouvirus mini-array 1 are identical to the repeats in the host CRISPR array. Thus, the pre-crRNA transcribed from this viral mini-array is predicted to be processed by the host CRISPR-Cas system and the mature viral crRNA would remain bound by the host effector complex^60^. Furthermore, the virus could potentially recruit the host CRISPR adaptation machinery to incorporate spacers into the mini-array and employ the respective crRNAs in inter-MGE competition, as demonstrated for some bacterial and archaeal micro-array carrying viruses^60, 64^, or for abrogation of host defenses. However, intriguingly, in the Chiyouvirus mini-array 1, we identified two self-targeting spacers, both with a 100% match to the corresponding protospacers (Fig. 3d). The role of these self-targeting spacers remains unclear. For the spacers in the Chiyouvirus mini-array 2, no potential targets were identified.

In addition to the mini-arrays, Chiyouvirus encodes a homolog of Cas2 nuclease (Chiyouvirus_39 in Extended Data Table S4), with a conserved Mg^2+^ binding site (Extended Data Fig. S9), which is an essential structural subunit of the adaptation complex in CRISPR-Cas systems. The virus-encoded Cas2 homolog might be a dominant negative inhibitor of spacer acquisition by the host CRISPR-Cas system.

Additionally, we attempted to predict Acrs among the Bathyarchaeia viral proteins by using a recently developed deep learning method^69^. We found that the structural model of a predicted Chiyouvirus Acr protein (Chiyouvirus_17 in Extended Data Table S4) was significantly similar to the N-terminal domain of AcrIF24 (7DTR, chain A) (Extended Data Fig. S10) which inhibits the activity of a type I-F CRISPR-Cas system^70^ and thus could function as an Acr as well.

Finally, in addition to the components of CRISPR-Cas systems, Fuxivirus and Kunpengvirus encode stand-alone DNA methyltransferases (Fuxivirus_47 and Kenpengvirus_43 in Extended Data Table S4) that could provide protection against the host restriction-modification systems.

To conclude, in this work, we identify four distinct groups of Bathyarchaeia viruses that can be expected to become new viral families. Notably, these viruses encode various components of CRISPR-Cas systems that could interfere with the host CRISPR-Cas immunity and/or mediate inter-virus conflicts. Thus, this study provides a glimpse into the virome of a widespread, comparatively abundant but poorly characterized class of archaea and the virus-host interactions in these organisms.

## Methods

### Bathyarchaeia genome collection and classification

Bathyarchaeia MAGs (metagenome-assembled genomes) were obtained from our previous dataset^8, 39^ and the NCBI (https://www.ncbi.nlm.nih.gov) and IMG/M (https://jgi.doe.gov) public database (up until June 30th, 2021). Bathyarchaeia genome classification was based on 51 high-coverage marker gene optimized from GTDB^71^ v207 (TIGR01171 and TIGR02389 were excluded due to the low coverage). Marker genes were initially identified with GTDB-Tk v2.1.1^72^ using ‘identify’ method. Only MAGs contain more than 50% of the marker genes were kept. Sequences were then aligned using MAFFT v7.457^73^ and trimmed with TrimAl v1.4^74^ (-gappyout). The maximum likelihood phylogenomic tree for concatenated 51 proteins was constructed with IQ-TREE 2.0.6 (best-fit model LG+F+R13, -B 1000, -alrt 1000). The quality, contamination, GC content, and other sequence information of Bathyarchaeia MAGs were assessed by checkM v1.2.2^75^. Biome information was extracted from the GenBank file or relevant literatures.

To delineate Bathyarchaeia orders, the maximum likelihood tree based on 51 proteins was converted to an ultra-metric tree and the relative evolutionary divergence (RED)^76^ was calculated. An order was called when its branch length in the ultrametric tree corresponded to a RED value within the range of 0.5 to 0.7.

### CRISPR-Cas system detection and CRISPR array validation

CRISPR-Cas systems in Bathyarchaeia MAGs were detected using DefenseFinder^77^ and further classified using CRISPRCasTyper^78^. CRISPR array in Bathyarchaeia MAGs were searched with MinCED^79^ (Default setting). To remove ambiguous CRISPR array that might not belong to Bathyarchaeia, all CRISPR array-containing contigs encoding proteins were searched against the nr database (2022-09) using Diamond (2.1.6) BLASTP^80^ (-evalue 1e-5, --more-sensitive), and only contigs with at least 3 proteins assigned to Bathyarchaeia were retained. The validated CRISPR repeats were clustered with 90% identity using an all-against-all BLASTN^81^ search (E-value cutoff 1e-5, word size 7).

### Bathyarchaeia virus identification and annotation

We first carried out the CRISPR-based virus-host assignments. Spacers from validated CRISPR array were used as a nucleotide database to compare to viral contigs in the IMG/VR v3^41^ database using BLASTN^81^ with parameters (-task blastn-short -evalue 1e-5). Only viral contigs harboring protospacers with 100% coverage and a maximum of one mismatch were considered as Bathyarchaeia viruses.

In addition, some viruses might form proviruses that cannot be detected by CRISPR- protospacer search. Therefore, we used VirSorter2 v.2.2.3^82^ to identified potential viral sequences in Bathyarchaeia MAGs. Specifically, all the Bathyarchaeia contigs were filtered using VirSorter2 (--include-groups dsDNAphage, ssDNA --min-length 5000 -- min-score 0.5) and only contigs with identifiable viral structural proteins and at least one ORF identified through a BLASTP search against the nr database as Bathyarchaeia were retained. Virial genome completeness estimation and the potential host regions trimming were conducted by CheckV 0.8.1^83^. Coding sequences were identified using Prokka^84^ with parameters (--kingdom viruses --gcode 1).

All the Bathyarchaeia viral proteins identified as describe above were annotated with DRAM^85^ (viral mode) and HHblits^86^ (MSA generated with UniRef30_2020_06 with 3 interactions, Evalue 1e-6, MSA was compared against the PDB_mmCIF70_14_Apr, SCOPe70_2.07, Pfam-A_v35 and UniProt-SwissProt-viral70_3_Nov_2021 databases with parameters -Z 250 -loc -z 1 -b 1 -B 250 -ssm 2 -sc 1 -seq 1 -dbstrlen 10000 - norealign -maxres 32000). All the viral genome maps were visualized using Proksee^87^.

### Bathyarchaeia viral taxonomy assignment

To determine the taxonomic status of Bathyarchaeia viruses, the whole genome network analysis was carried out with vConTACT2^88^ (default parameters) against the Viral RefSeq v207 database as well as archaeal viruses proposed in the latest International Committee on Taxonomy of Viruses Report (VMR_MSL38_v1). The resulting networks were visualized using Cytoscape v3.9.1^89^ with an edge-weighted spring embedded model.

For viruses in the realm *Duplodnaviria*, proteome-scale phylogeny was constructed using the VipTree version 3.4^90^. The analysis was carried out with archaea viral families in the realm *Duplodnaviria*, according to the latest ICTV taxonomy classification (04/25/2023). For filamentous viruses, proteome-scale phylogeny was constructed by using VICTOR^91^ with all known members of the *Tokiviricetes* class. Structural modeling of viral proteins was performed with AlphaFold2^92^ via ColabFold v1.5.1^93^ with pdb70 template mode.

### Bathyarchaeia viral CRISPR-Cas systems classification

All the Cas proteins were manually search with HHblits (see above). To determine the CRISPR-Cas system subtype of ‘*Fuxiviridae*’, phylogenetic tree of all the Cas proteins were constructed with reference sequences. Specifically, Cas4 proteins were aligned with 29 reference^67^ sequences by MUSCLE 5.1^94^ and trimmed by TrimAl v1.4 (-gappyout). The maximum likelihood tree was inferred by IQ-TREE 2.0.6 (-m LG+I+G4, -B 1000, -alrt 1000). Cas7 proteins were aligned with 296 reference sequences^95^ by MUSCLE 5.1. The alignment was trimmed by TrimAl v1.4 (-gt 0.7) and sequence with more than 50% gaps were dropped. The maximum likelihood tree was inferred by IQ-TREE 2.0.6 (-m VT+R10, -B 1000 -alrt 1000). Cas5 proteins were aligned with 722 reference^95^ sequences by MUSCLE 5.1. The alignment was trimmed by TrimAl v1.4 (-gappyout) and sequence with more than 50% gaps were dropped. The maximum likelihood tree was inferred by IQ-TREE 2.0.6 (-m VT+R10, -B 1000 -alrt 1000). Cas11 proteins were aligned with 45 reference sequences^59, 95^ by MUSCLE 5.1. The alignment was trimmed by TrimAl v1.4 (-gappyout) and sequence with more than 50% gaps were dropped. The maximum likelihood tree was inferred by IQ-TREE 2.0.6 (-m VT+R3, -B 1000 -alrt 1000). Csf1 proteins were aligned with 49 reference sequences^59, 95^ sequences by MUSCLE 5.1. The alignment was trimmed by TrimAl v1.4 (-gappyout) and sequence with more than 50% gaps were dropped. The maximum likelihood tree was inferred by IQ-TREE 2.0.6 (-m WAG+I+G4, -B 1000 -alrt 1000). Phosphoadenosine phosphosulfate (PAPS) reductase (CysH) domain containing protein has been observed in prokaryotes and virus^61^. We first recalled all the CysH-like proteins from 18988 public reference complete viral genomes^96^. Together with *bona fide* CysH and Cas-associated CysH-like proteins, 322 sequences were aligned by MUSCLE 5.1 and trimmed by TrimAl v1.4 (-gappyout). The maximum likelihood tree was inferred by IQ-TREE 2.0.6 (-m WAG+F+R7, -B 1000 -alrt 1000).

All final phylogenetic trees were visualized by tvBOT^97^.

### Bathyarchaeia viral CRISPR spacer-protospacer match analysis

CRISPR array in Bathyarchaeia viral genome were detected with MinCED (-minNR 1). All the spacer obtained from Bathyarchaeia virus were BLASTN search (-task blastn-short) with viral database IMG/VR v3 and plasmid sequences database PLSDB database^98^ (v. 2021_06_23_v2) as well as Bathyarchaeia MAGs in this study. Direct repeats were searched against CRISPR-CAS++ databases^99^ and Bathyarchaeia CRISPR repeat dataset with BLASTN (-evalue 0.01).

### Bathyarchaeia viral anti-CRISPRs proteins (Acrs) identification

The potential anti-CRISPR proteins (Acrs) of Bathyarchaeia viruses were first predicted using DeepAcr^69^. The protein structures of predicted Acrs, along with Acrs from reference^100^, were modeled using AlphaFold2 by ColabFold v1.5.2 in pdb70 template mode. These potential Bathyarchaeia virus Acrs were then used as query models for alignment to the reference Acr models using TM-align^101^, and only models with a TM score greater than 0.3 were retained. This initial computational analysis was further supplemented with comprehensive meticulous manual inspection of the alignments and structural models.

### Etymology

‘*Fuxiviridae*’: Derived from Fuxi, a legendary figure in Chinese mythology known for his diverse talents and abilities. This alludes to the possibility that the virus has a multifaceted counter-defense systems, capable of employing various strategies to evade the host’s immune response.

‘*Kunpengviridae*’: Named after Kunpeng, a mythical creature in Chinese mythology known for its transformative abilities. The name alludes to the virus’s ability to integrate into the host genome to form proviruses.

‘*Huangdiviridae*’: Named after Huangdi, the legendary Chinese sovereign often associated with important inventions. Given Huangdi’s connections (the host MAG of Huangdivirus in the Baizomonadales order) in Chinese mythology.

‘*Chiyouviridae*’: Inspired by Chiyou, a symbol of war and invention in Chinese mythology.

## Acknowledgments

We would like to express our deepest gratitude to the following professors for their generous support and data sharing, which has been invaluable to our research. Special thanks go to Professors Natasha Ivanova, Robert Kelly, Tanja Woyke, William P. Inskeep, Jim Fredrickson, Roland Hatzenpichler, Brian P. Hedlund, and Ramunas Stepanauskas for their enthusiastic assistance and valuable suggestions. We sincerely appreciate their contributions. Also, thanks to Dr. Jie Pan, Mr. Chengxiang Gu and Mr. Jinquan Li for their invaluable guidance and assistance.

This work was supported by grants from the National Key Research and Development Program of China (2022YFA0912200), the National Natural Science Foundation of China (32225003, 92251306,31970105), the Innovation Team Project of Universities in Guangdong Province (No. 2020KCXTD023), the Shenzhen Science and Technology Program (JCYJ20200109105010363), and Shenzhen University 2035 Program for Excellent Research (2022B002). E.V.K. is supported by the Intramural Research Program of the National Insitutes of Health of the USA (National Library of Medicine).

## Author contributions

M.L. and C.D. designed the experiments; C.D., Yang L., Ying L. and M.K. provided support on bioinformatic analysis. C.D. performed the data analyses and wrote the paper with contributions from all the authors.

## Extended Data

Extended Data Fig. S1: The calculated relative evolutionary divergence (RED) value of each node at order level. Eight order-level units of class Bathyarchaeia are marked in different colors.

Extended Data Fig. S2: Bathyarchaeia CRISPR Array Overview. a) CRISPR repeats found in Bathyarchaeia MAGs, clustered at 90% sequence identity. Verified repeats are highlight in green, while repeats associated with detected viruses in this study are indicated in red. b) Seq-logo for repeat clusters linked to viruses identified in this research. c) Bathyarchaeia CRISPR spacer length distribution.

Extended Data Fig. S3: The genome-wide sequence similarity comparison, phylogenetic and modeling of major capsid of Bathyarchaeia viruses in realm *Duplodnaviria*. a) The proteomic tree displays the relationship between Bathyarchaeia viruses and archaea viruses in the realm *Duplodnaviria*, based on genome-wide sequence similarities. Bathyarchaeia virus families are labeled in green, and viruses with complete genomes are marked with a red circle. b) The maximum likelihood tree of major capsid proteins (MCPs) shows the relationship between Bathyarchaeia virus and archaeal virus. Bathyarchaeia virus families are labeled in green, and the 3D structure representing the MCP of each family is displayed next to it.

Extended Data Fig. S4: Reads coverage of Fuxivirus genome. Raw reads from data (SRR5248875) are mapped to the virus genome with 100% identity using Bowtie 2 and Samtools v1.1.7.

Extended Data Fig. S5: Huangdivirus structural proteins. a) Structural modeling of Huangdivirus capsid proteins VP1 and VP3, colored using a rainbow gradient from N-terminus (blue) to C-terminus (red). b) Sequence analysis of Huangdivirus structural proteins VP1-3, with predicted transmembrane domains highlighted in yellow and theoretical glycosylation consensus motifs (N-X-S/T) shown on a green background.

Extended Data Fig. S6: Genomic analysis of Chiyouvirus. a) Phylogenomic tree of Chiyouvirus (red arrow) alongside known members of the *Tokiviricetes* class, based on whole-genome amino acid analysis using VICTOR. Tree is rooted with *Primavirales*, and branch length represents GBDP distance formula D6. Branch support values are indicated with numbers. b) Whole-genome amino acid identity comparison of filamentous viruses in the *Tokiviricetes* class, conducted by EzAAI^104^. Chiyouvirus is highlighted in red. Only AAI values greater than 40% are displayed in the heatmap. c) Predicted structural model comparison of Bathyarchaeial Chiyouvirus major capsid proteins MCP1 and MCP2 with Icerudivirus SIRV (3J9X, chain A) and Captovirus AFV1 (5W7G, chain A) structures. Models are colored using a rainbow gradient from N-terminus (blue) to C-terminus (red).

Extended Data Fig. S7: Phylogenetic analysis of proteins in Fuxivirus type IV-B CRISPR-Cas system and Kunpengvirus Cas4 protein. Maximum likelihood trees for a) Csf2 (Cas7), b) Cas4 c) CysH-like protein, d) Csf3 (Cas5), e) Csf1, and f) Csf4 (Cas11). Fuxivirus proteins are marked with red pentagrams. Kunpengvirus Cas4 protein is highlighted with blue in figure b).

Extended Data Fig. S8: Sequences alignment of Cas4 protein. Conserved amino acid residues involved in the functions of CRISPR spacer acquisition are highlight in red dot.

Extended Data Fig. S9: Sequences alignment of Cas2 protein. Conserved amino acid residues involved in Mg^2+^ binding site is highlight in red dot.

Extended Data Fig. S10: Structure alignment of predicted anti-CRISPR protein (Acr) of Chiyouvirus with reference AcrIF24. Models of Bathyarchaeia virus encoding Acr is colored in green, AcrIF24 NTD is colored in wheat. AcrIF24 CTD is colored in palegcyan.

**Table S1. Detailed statistics of Bathyarchaeia MAGs.**

**Table S2. Viral contigs targeted by Bathyarchaeia spacers.**

**Table S3. Classification of Bathyarchaeia viruses.**

**Table S4. Annotations of proteins encoded by Bathyarchaeia viruses by DRAM-v and HHblits.**

**Table S5. Comparison of Bathyarchaeia virus encoding protein with other archaea viruses.**

