## Supplementary figures and images for "Diversity of Bathyarchaeia viruses in metagenomes and virus-encoded CRISPR system components" (from another version of this preprint: version 1, 2023-08-24)

### Supplemental Fig. S1

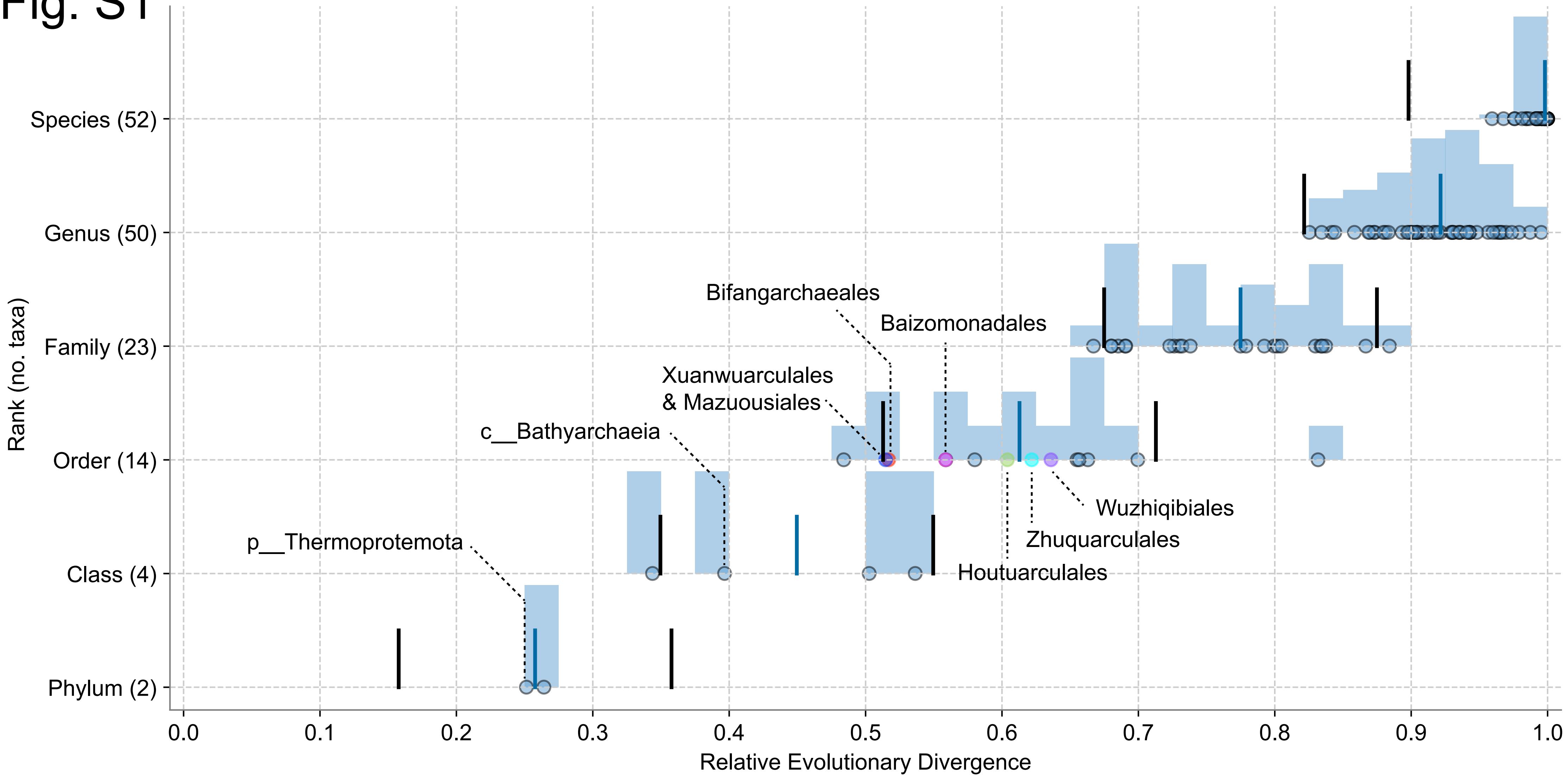

### Supplemental Fig. S2

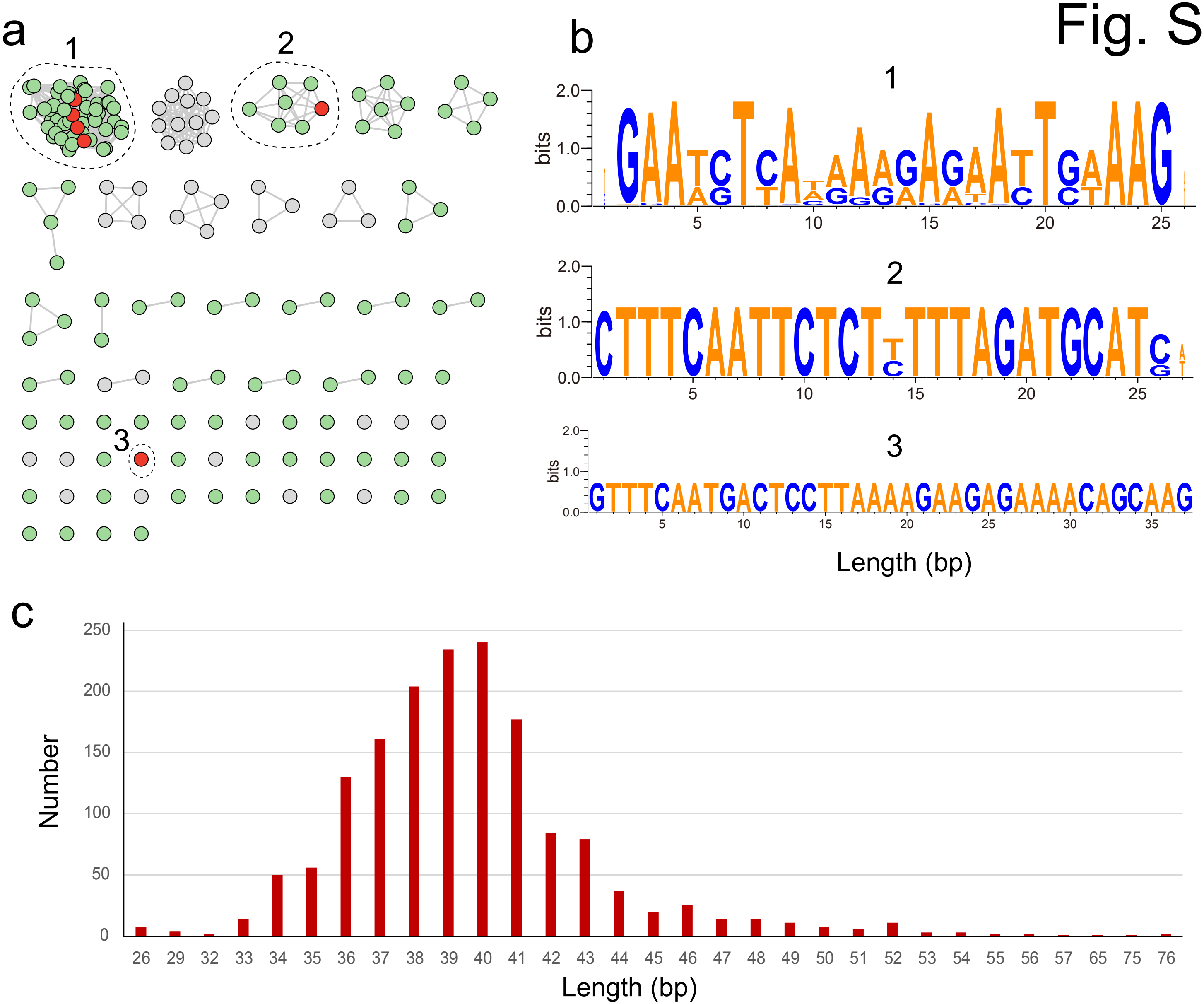

### Supplemental Fig. S3

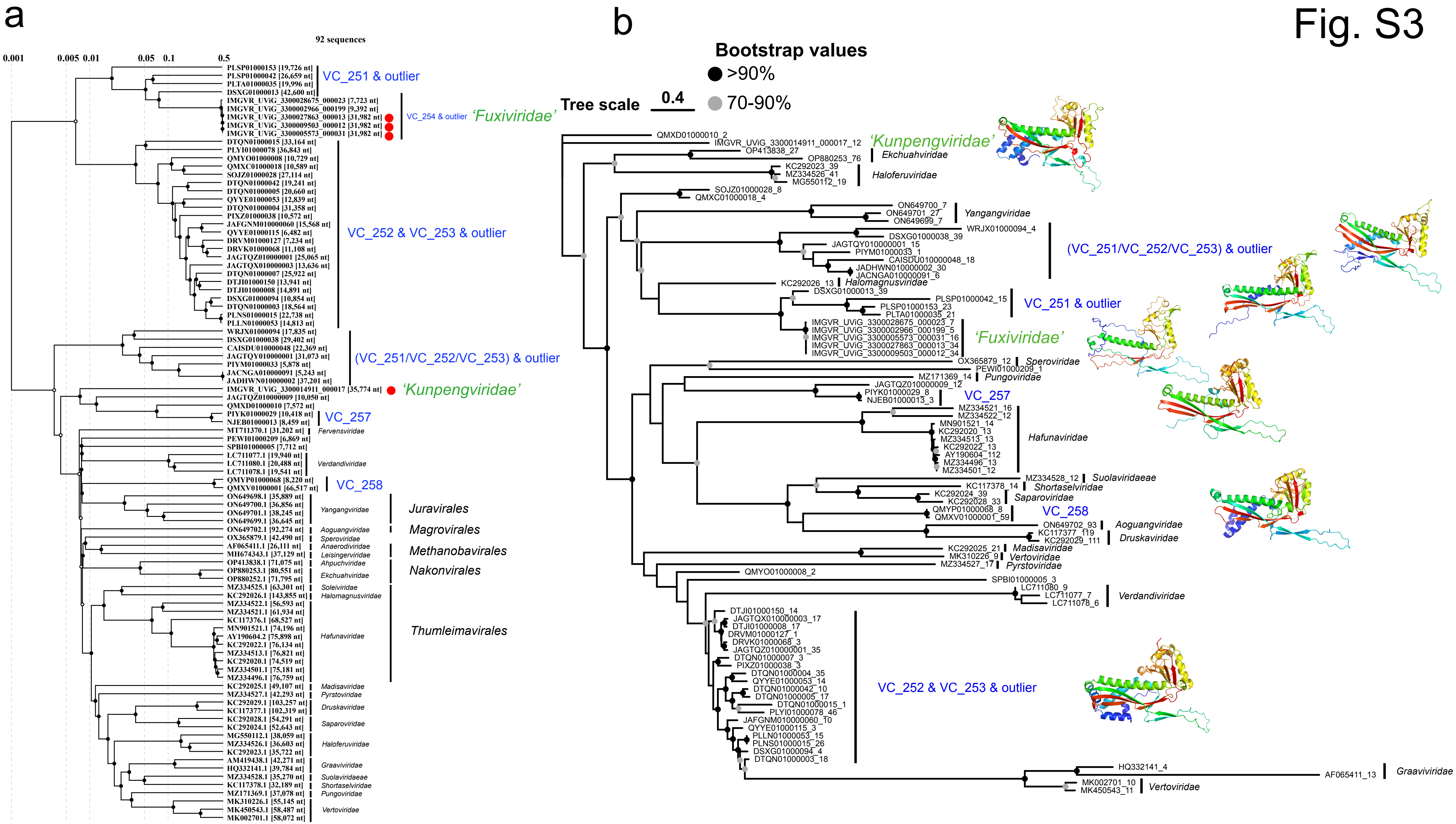

### Supplemental Fig. S4

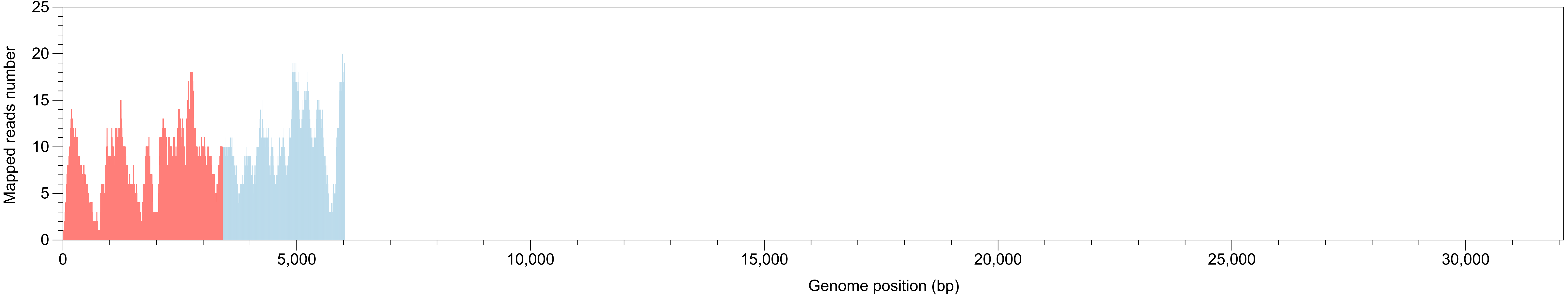

### Supplemental Fig. S6

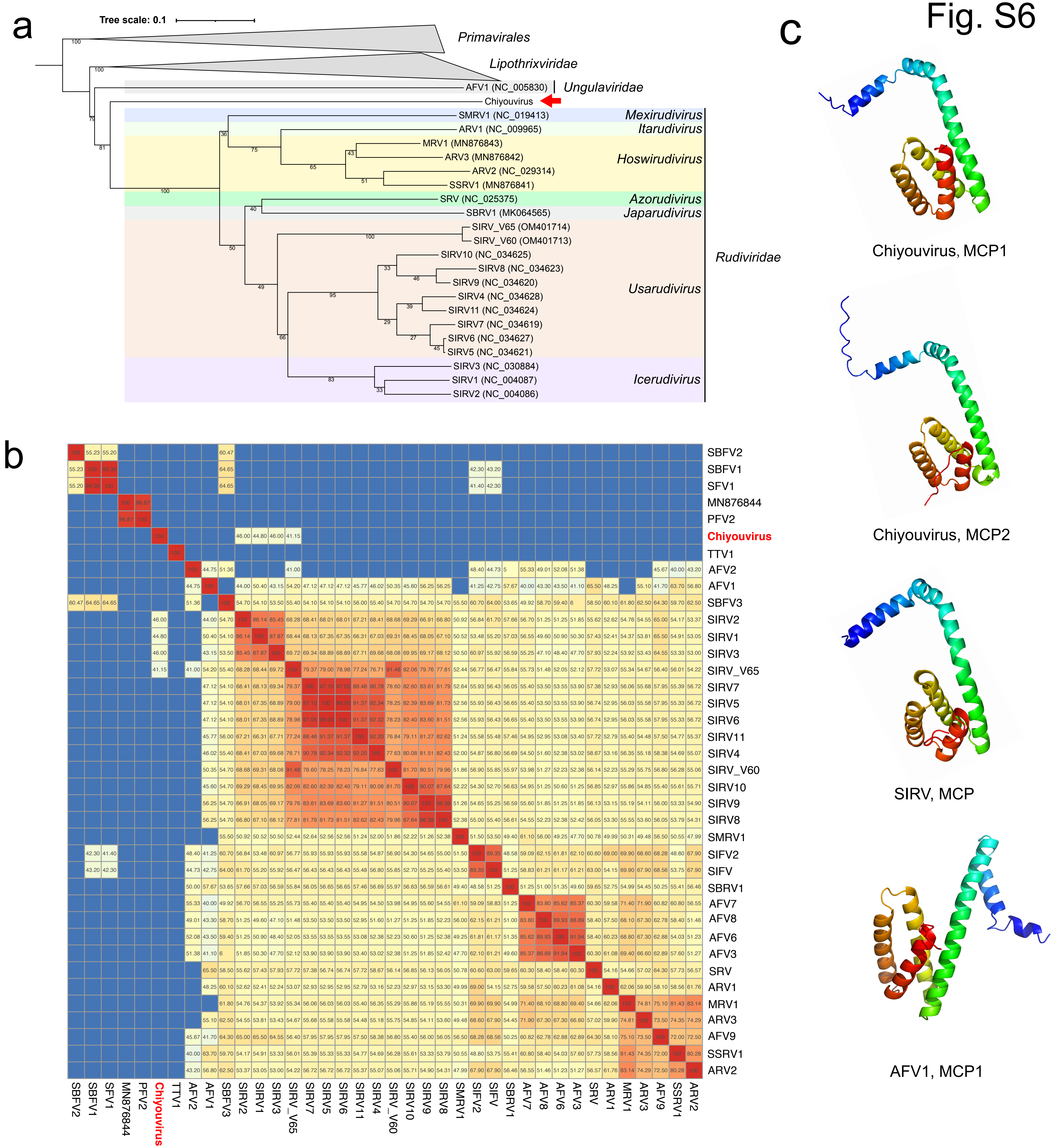

### Supplemental Fig. S7

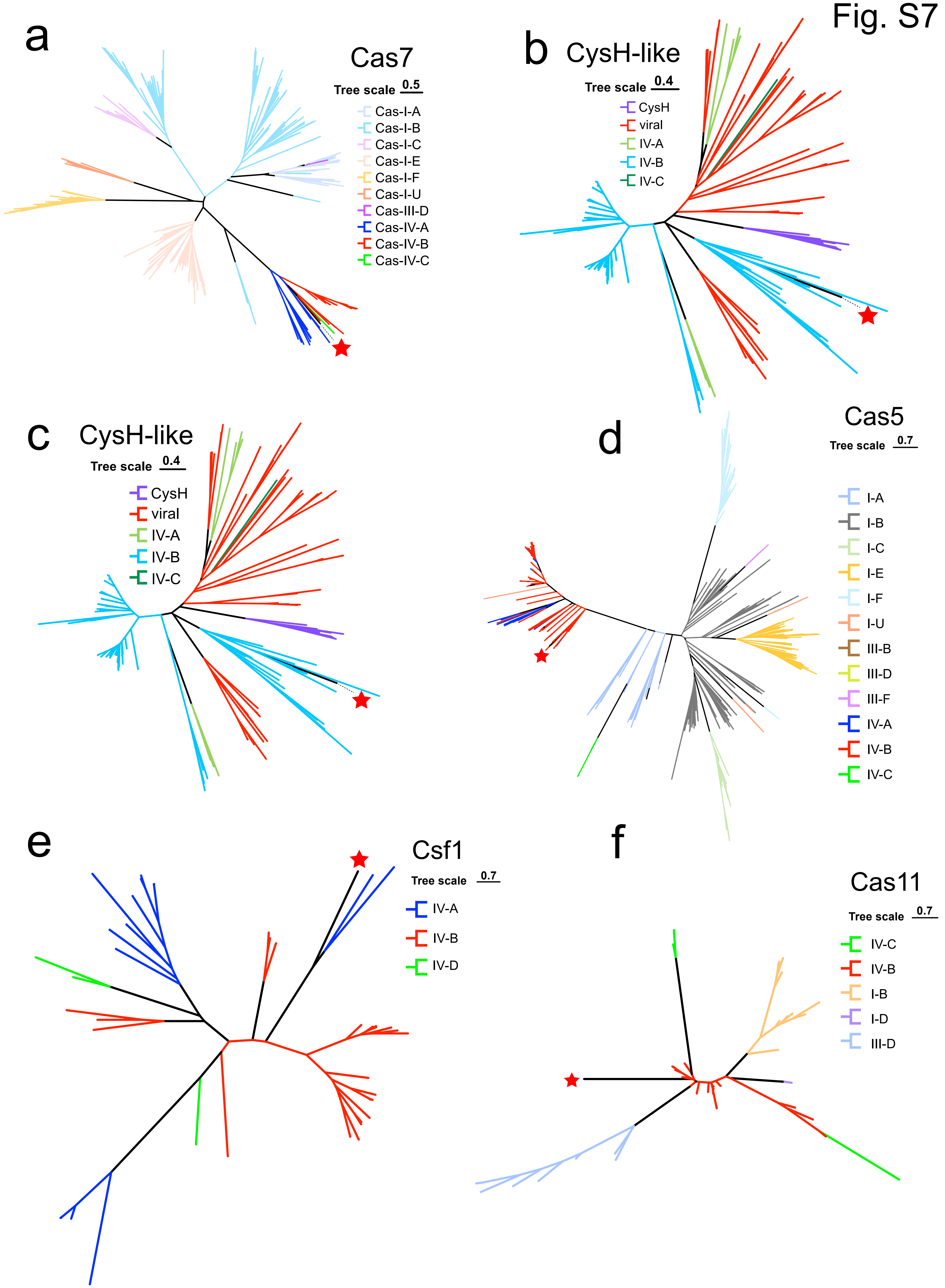

### Supplemental Fig. S8

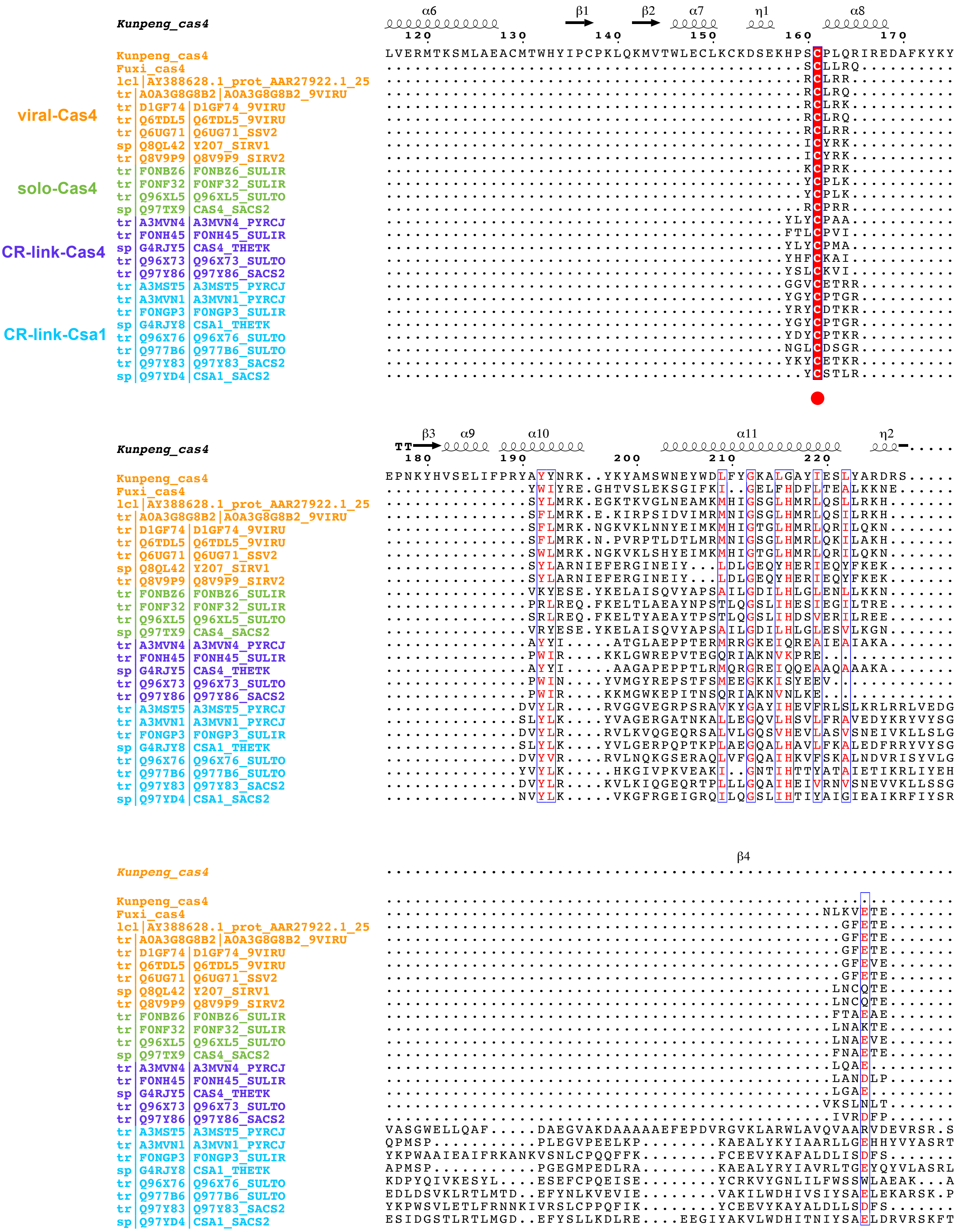

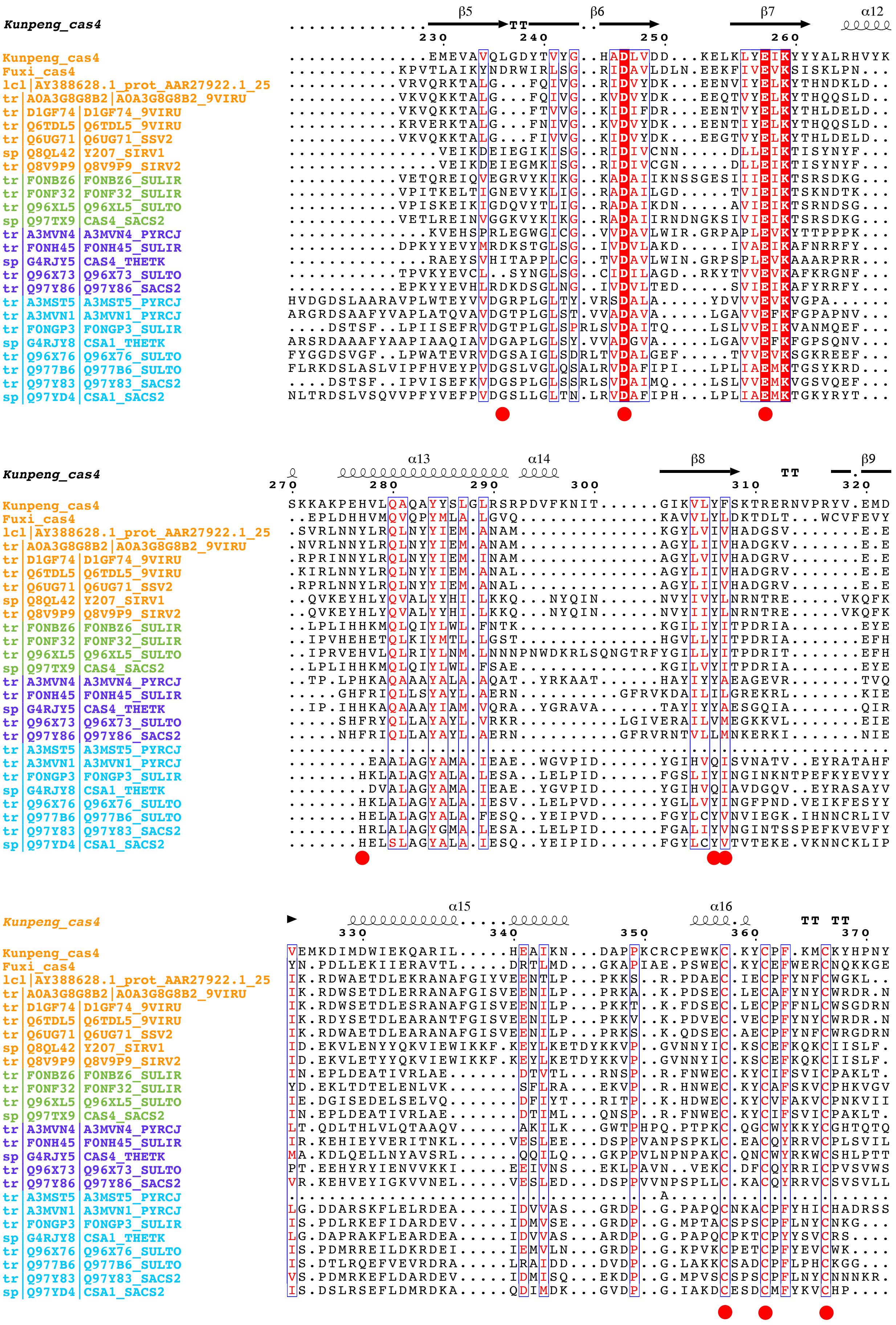
