## Supplemental Fig. S5 for "Diversity of Bathyarchaeia viruses in metagenomes and virus-encoded CRISPR system components" (from another version of this preprint: version 1, 2023-08-24)

a

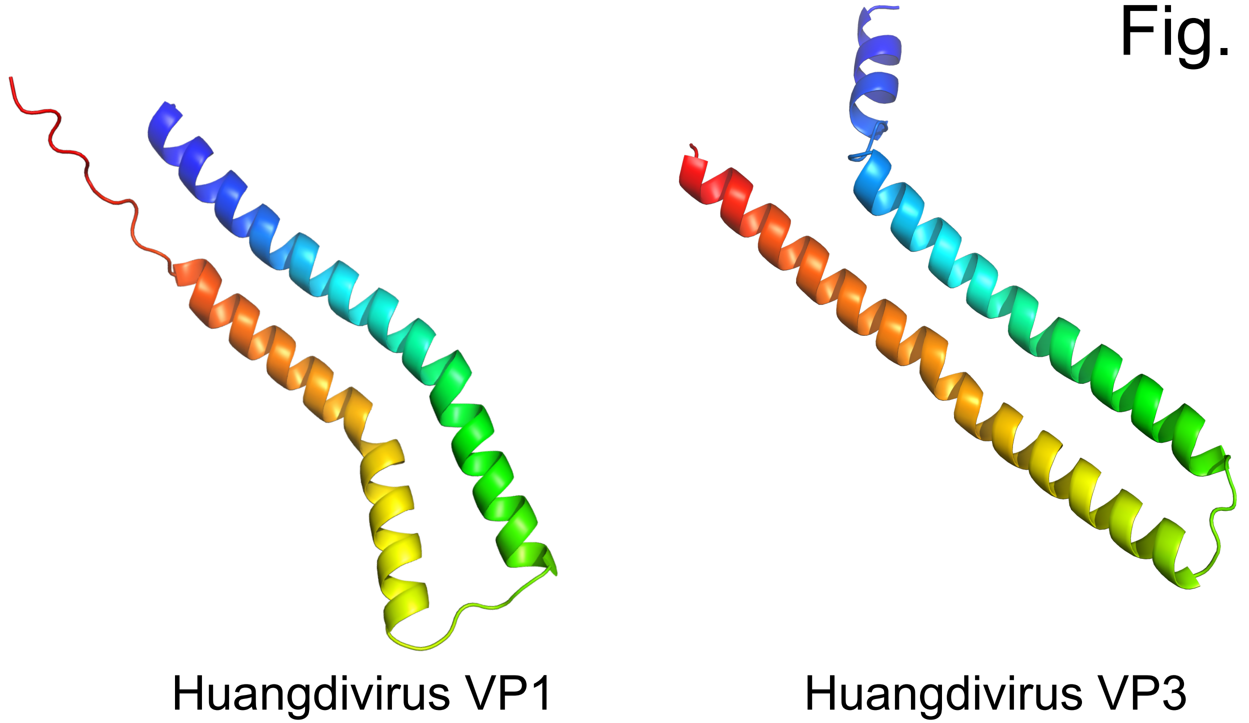


b

```
                α1                                                     
VP1_structure
            1        10        20        30        40        50        60
VP1         M.......KSLRRNRSGLIYVWVVCFFAIVLYSIVWFVLGWPAMMTIQAVEDAYTFTGPAATTVDLV
VP3         MEMLAKLLKNVRGDSSVIALVIMLILFGVVL..PVGLMVTGSLYTTVSNMDLGSQGNSTRTTLFNNI

                α2
VP1_structure
                      70        80        90
VP1         KTVIAWH...PLIFIFGMIIWALVNSHKREYVSYQEG.
VP3         YSAYNLSVILPIVAAAGMIIAAIM.....LYFRWRSST

            1        10        20        30        40        50        60
VP2         MAKKEKWIQKVVKPNERGLLHKQLGISTKERIPKTLVQKIVKTPIGKKIQNPTKTGKRTI
                     70        80
VP2         KVTRLLKQRANFALNVGYGRYPRRK
```
