## Supplemental Fig. S9 for "Diversity of Bathyarchaeia viruses in metagenomes and virus-encoded CRISPR system components" (from another version of this preprint: version 1, 2023-08-24)

```
                                                                  β1         TT    TT       α1
Chiyouvirus1_Cas2

Chiyouvirus1_Cas2       1   ...........................MYLVVMDKPGA.KPGRMPRTFYYRLKKMNLPVT
sp|Q9KFX8|CAS2_HALH5    1   ..........................MLVLITYDVQTSSMGGTKRLRKVAK.ACQNYGQ.
sp|Q97YC2|CAS2A_SACS2   1   MA........................MLYLIFYDITDD.....NLRNRVAE.FLKKKGLD
sp|A0Q5Y5|CAS2A_FRATN   1   MQ.LR.....................KEYLIAYDIEDN.....KTRTIIYK.QLLAYGLK
sp|C9X1G7|CAS2_NEIM8    1   MSEAK....................FMRIVFFDLPVITAAKRKAANQFRQ.FLLKDGYQ
sp|D4GQP1|CAS2_HALVD    1   ..........................MQVIVVYDVPAK.......RTRIYR.KLLRRRLE
sp|G3ECR3|CAS2_STRTR    1   MS.YR....................YMRMILMFDMPTDTAEERKAYRKFRK.FLLSEGFI
sp|P9WPJ3|CAS2_MYCTU    1   MP.TRSREEYFNLPLKVDESSGTIGKMFVLVIYDISDN....RRRASLAK.ILAGFGY.
```

```
                            β2  TT  β3                α2                        β4
Chiyouvirus1_Cas2

Chiyouvirus1_Cas2      33   RVQKSVYVTES......RSAAYRLAGLARE....YGFMVKVY..KVVEE...........
sp|Q9KFX8|CAS2_HALH5   33   RVQNSVFECIV.DSTQLTSLKLELTSLIDE....EKDSLRIY.......RLGNNYKTKVE
sp|Q97YC2|CAS2A_SACS2  31   RIQYSVFMGDL.NSSRLKDVEAGLKIIGNRKKLQEDERFFILIVPITENQFRE..RIVIG
sp|A0Q5Y5|CAS2A_FRATN  33   AVQKSVFWGYV.SIAELNAIKRLFDSSLT.....ISDKVFITRVNMHEQKLDYSFGYDDK
sp|C9X1G7|CAS2_NEIM8   40   MLQLSVYSRIVKGRDSLQKHHNRLCANLP.....QEGSIRCL..EITEKQY....AAMKL
sp|D4GQP1|CAS2_HALVD   27   HLQYSVFFGEL.TAGQVTAMKNEIESELE.....PTDSIVVF.......EFDNPHAFDHT
sp|G3ECR3|CAS2_STRTR   39   MHQFSVYSKLLLNHTANTAMVGRLKANNP.....KKGNITIL..TVTEKQF....ARMIY
sp|P9WPJ3|CAS2_MYCTU   53   RVQESAFEAML.TKGQLAKLVARIDRFAI.....DCDNIRIY.......KIRG..VAAVT
```

```
Chiyouvirus1_Cas2

Chiyouvirus1_Cas2           .............................
sp|Q9KFX8|CAS2_HALH5   81   HIGAKP....SIDLEDPLIF.........
sp|Q97YC2|CAS2A_SACS2  88   YSGS........EREEKSNVVW.......
sp|A0Q5Y5|CAS2A_FRATN  87   TFK.........DWDEYGHI.........
sp|C9X1G7|CAS2_NEIM8   89   LLGELKTQEKKVNSDQLLLF.........
sp|D4GQP1|CAS2_HALVD   74   TFG.........DADEPGSRFT.......
sp|G3ECR3|CAS2_STRTR   88   LYGDKNTS..IANSEERLVFLGDNYCDED
sp|P9WPJ3|CAS2_MYCTU   98   FYGRGR....LVSAEEFVFF.........
```
